# Measurement intensification for antibody formulations: combined measurement of protein size, interactions, and viscosity by differential dynamic microscopy

**DOI:** 10.64898/2026.01.20.700608

**Authors:** Caidric I. Gupit, Anoushka Shandilya, Juan Manuel Urueña, Nadya Morales-Cummings, Rohini Gupta, Megan T. Valentine, Matthew E. Helgeson

## Abstract

High-throughput screening and optimization of high-value protein formulations requires intensified measurements to extract a wide range of properties using a small number of measurement techniques, small sample volumes, and short measurement times. We demonstrate how differential dynamic microscopy (DDM) can fill this need by measuring a broad range of key biophysical properties relevant to protein formulations from a single workflow on microliter-scale samples using label-free video optical microscopy. We show that the use of phase contrast imaging dramatically enhances measurement resolution for protein solutions at dilute and semidilute concentrations, enabling measurement of colloidal properties such as protein-protein interactions, protein size, aggregation, and solution viscosity from a single set of measurements. DDM measurements on a representative human immunoglobulin (IgG) system yield estimates for the hydrodynamic radius (*R*_h_), second osmotic virial coefficient (*B*_2_), and hydrodynamic interaction (*k*_d_) that are consistent with independently measured values, validating the ability of DDM to extract these parameters from a single set of measurements. Observed trends in *B*_2_ with pH and ionic strength are consistent with the antibody’s charge and screened electrostatics, demonstrating the ability of DDM to provide insight on protein-protein interactions. To show the utility of DDM as a “multitool” for quantifying multiple formulation properties from a single measurement, we use the results to test a predictive colloidal model for the solution viscosity, which is in fair agreement with measurements obtained using DDM-based microrheology. Combined with low sample requirements and short measurement times, DDM thus offers a high-throughput and efficient route to accelerate protein biophysics and formulation development.

**SIGNIFICANCE STATEMENT:** The formulation of stable, high-concentration antibodies and other protein solutions requires extensive biophysical measurements that are often material- and time-intensive. We demonstrate that differential dynamic microscopy (DDM) provides a powerful alternative by providing rapid access to a broad range of industrially relevant colloidal properties from a single measurement on microliter-scale samples using conventional video optical microscopy. This capability makes DDM an attractive, low-resource approach for routine biomolecular formulation screening and optimization.

## INTRODUCTION

Antibody-based biopharmaceuticals constitute a multi-billion-dollar global industry that drives the development, formulation, and approval of new antibody therapeutics for diseases ranging from cancer and infectious diseases to autoimmune disorders (1). As subcutaneous and intramuscular delivery become more common, there is an increasing demand for highly concentrated formulations (>50 mg/mL) to enable effective dosing within the restricted injection volumes (<2 mL) available for these routes. However, such high concentrations often result in increased viscosity and protein aggregation,(2, 3) which present significant challenges in formulation development, manufacturing, stability, and administration. For example, high viscosity can necessitate prohibitively strong injection forces to deliver the formulation through a needle, while antibody aggregates can compromise efficacy and trigger adverse immune responses in patients.

Due to the simultaneous constraints of rapid development and concentrated formulations, predicting and avoiding these compromising effects is crucial in the early stages of antibody product development. However, doing so necessitates characterizing a range of biophysical properties to understand, control, and predict the colloidal properties of antibodies within a formulation. This includes measurements of protein-protein interactions, protein size, aggregation propensity, and formulation viscosity. Conventionally, each of these properties is determined using different well-established measurement techniques and corresponding instrumentation such as analytical ultracentrifugation,(4) static(5) and dynamic(6) light scattering, rheometry,(7) and various chromatographic methods(8–10). The need for multiple techniques dramatically increases the material and time required for formulation screening and optimization, creating a significant bottleneck in the development of new biopharmaceutical products(2). Consequently, there is an increased demand for biophysical characterization methods that can provide access to a wide range of properties from fewer measurements, with reduced sample volumes and measurement times.

One potential strategy to circumvent the need for diverse biophysical measurements is the development of accurate predictive models and correlations. For example, previous studies proposed that increased viscosity and aggregation in high-concentration antibody formulations can be correlated with low-resolution measures of nonspecific protein-protein interactions at lower concentrations, such as the second osmotic virial coefficient, *B*_2_,(4, 11) or the diffusion interaction parameter, *k*_d_(4, 7, 12–14). However, these correlations are highly dependent on the specific details of the formulation,(7, 15, 16) and therefore must be developed individually, limiting their general applicability. Data-driven models have shown some success in correlating viscosity with antibody sequence(17), although the transferability of these models across a wide range of formulations has yet to be proven. More broadly-applicable theoretical models require biophysical measurements of colloidal properties including protein-protein interactions and hydrodynamic interactions, which are typically accessible only through more advanced and resource-intensive techniques such as X-ray and neutron scattering (18).

An alternative approach involves the intensification and miniaturization of antibody biophysical property characterization into multi-modal measurements that provide a wealth of information from a single or small number of low-volume samples. Optical microscopy is particularly well-suited for this purpose due to its relatively low cost, simple instrumentation, low material requirements, and straightforward integration with automation. In particular, differential dynamic microscopy (DDM), which combines simple video microscopy with Fourier-based image analysis reminiscent of light scattering,(19, 20) has emerged as a powerful technique for multi-modal analysis of complex fluid structure and dynamics(21, 22). For example, DDM has been successfully applied to characterize a wide range of dynamic phenomena, from the Brownian dynamics and particle sizing of diluted(23) and concentrated(24) colloidal suspensions, diffusive dynamics of polydisperse protein-rich clusters,(25) adsorption of proteins onto nanoparticles,(26) roto-translational dynamics of anisotropic samples,(27) constrained motion in crowded systems,(28) kinetics of polymerization,(29) gelation,(30) polyelectrolyte condensates,(31) and active motion of microorganisms(32) and cells(33).

Due to its ability to extract a wide range of colloidal properties of bio- and macromolecules without requiring external probes or complex labeling schemes while using minimal sample volumes, we hypothesize that DDM is ideally suited as a high-throughput and potentially automatable method for intensified protein formulation characterization and optimization. Previous studies successfully demonstrated the use of DDM for characterizing protein size and aggregation (25, 34). More recently, its application has been extended to perform microrheology measurements,(29, 35) including high-throughput, fully autonomous DDM microrheology for optimizing the viscosity and gelation of protein formulations(30, 36). Of particular relevance to the present work, DDM has also been demonstrated as an alternative to conventional static light scattering to estimate *B*_2_ from concentration-dependent measurements of protein solutions(34). These studies show that DDM measurements can be performed using relatively unsophisticated bright-field microscopy, without the need for fluorescent labeling or colloidal probes that might interfere with the properties or stability of the protein formulation. However, these measurements were primarily focused on globular or intrinsically disordered proteins with isotropic shape and relatively simple protein-protein interactions. Consequently, the applicability of DDM to the characterize antibodies, which have an anisotropic shape and complex interactions, has yet to be assessed. More broadly, studies that integrate the determination of all these diverse properties using DDM from a single small set of measurements would better demonstrate its potential as a multiplexed tool for the biophysical characterization of biotherapeutic formulations.

In this work, we demonstrate the versatility of DDM to fill this role for characterizing antibody formulations through measurements of *B*_2_and *k*_d_, protein size and aggregation stability, as well as solution viscosity from a single experimental workflow (**Figure 1**). Viscosity was determined via DDM microrheology by analyzing the dynamics of added inert probe particles. The remaining properties were all obtained from a single concentration series of probe-free samples by detecting sub-diffraction-limit intensity fluctuations using phase-contrast imaging, which increases the signal-to-noise ratio by nearly an order of magnitude compared with standard bright-field imaging. The measured quantities were validated against results from corresponding conventional biophysical techniques, and the results were combined to test a previously proposed predictive theory for the viscosity of protein solutions from their colloidal properties(18). By providing rapid access to a broad range of industrially relevant biophysical properties with minimal sample requirements and simple inexpensive instrumentation, DDM offers a powerful approach for developing stable antibody formulations in biopharmaceutical applications. Its compatibility with fully automated, unsupervised, and high-throughput workflows(30) further enhances its appeal as a tool for routine biomolecular formulation screening and optimization.

**Figure 1.**
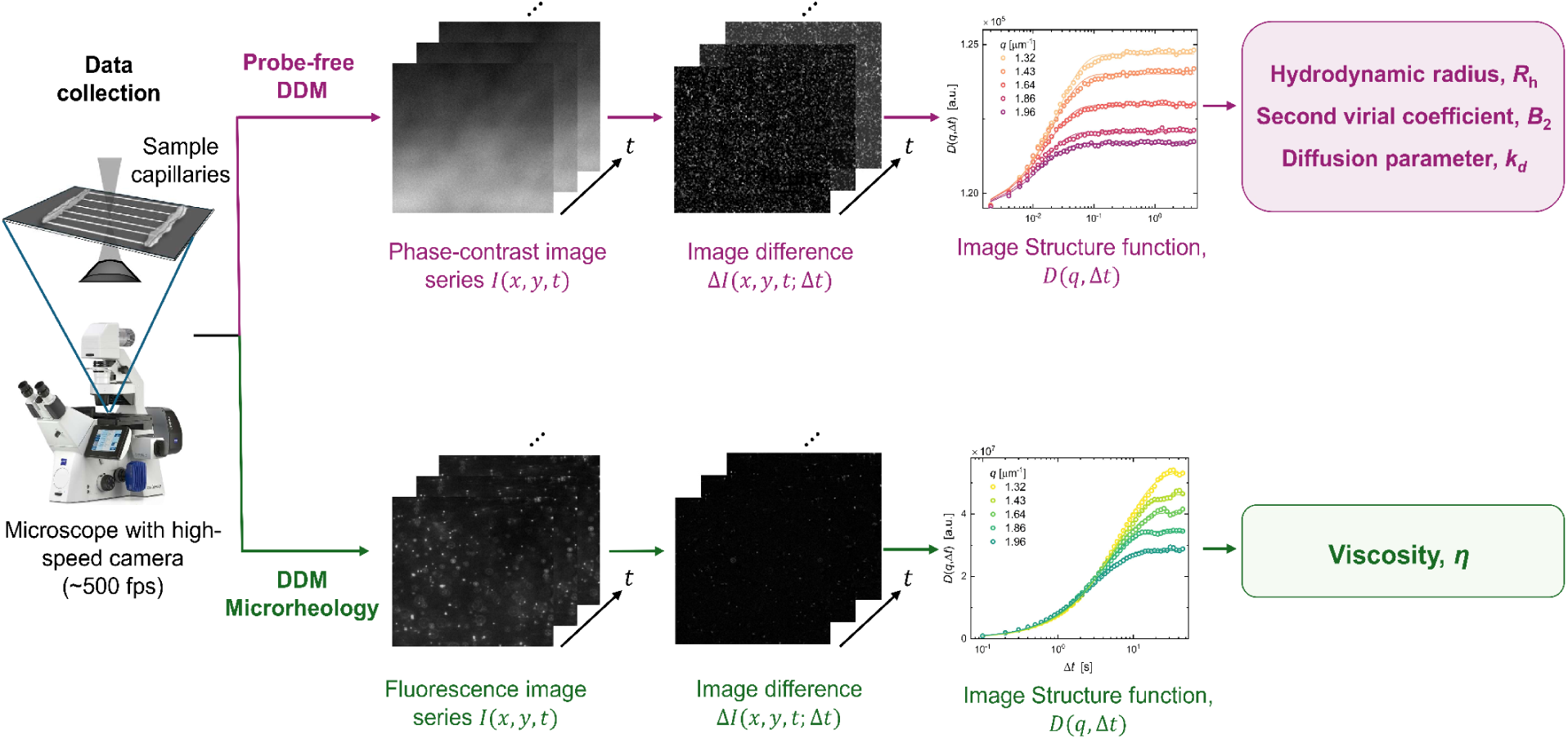
Schematic representation of probe-free and microrheology DDM workflows. In probe-free DDM (top row), phase-contrast image series *I*(*x*, *y*, *t*) are acquired and processed into image differences Δ*I*(*x*, *y*, *t*; Δ*t*), from which the image structure function *D*(*q*, Δ*t*) is computed to extract the hydrodynamic radius *R*_h_, second virial coefficient *B*_2_, and diffusion parameter *k*_d_. In microrheology DDM (bottom row), fluorescence image series *I*(*x*, *y*, *t*) of probe particles are similarly analyzed to obtain *D*(*q*, Δ*t*) and determine the sample viscosity *η*.

## 1. MATERIALS AND METHODS

### 1.1. Materials

The chemicals used in this work were purchased from Sigma-Aldrich (St. Louis, MO, USA) and used without further purification. The test antibody used was polyclonal Immunoglobulin G (IgG from human serum). The polyclonal nature of the protein sample leads to a heterogeneous distribution of its biophysical properties, which will become important in the interpretation of the experimental results to follow. Sodium acetate, glacial acetic acid, 2-(*N*-morpholino)ethanesulfonic acid (MES) monohydrate, tris(hydroxymethyl)aminomethane (Tris), L-histidine, L-histidine dihydrochloride, sodium chloride, sodium hydroxide, and hydrochloric acid were all used as received.

### 1.2. Sample Preparation

Sodium acetate (25 mM, pH 5), MES (25 mM, pH 6), Tris (25 mM, pH 7, 8, 9), and histidine (20 mM, pH 6) buffers were prepared in Milli-Q water and adjusted to the target pH using a calibrated pH meter (SevenDirect SD50; Mettler-Toledo, Columbus, OH, USA). Two series of samples were prepared, one with varying pH using sodium acetate buffer, MES buffer, and Tris buffer to cover a pH range from 5 to 9 with added 100 mM NaCl, and the other with varying concentration of added NaCl from 0 to 200 mM using histidine buffer at pH 6. IgG solids were dissolved in the appropriate buffer and serially diluted to prepare samples at a series of IgG concentrations ranging from 0.8 to 82.2 mg/mL. Protein LoBind^®^ microcentrifuge tubes (Eppendorf, Hamburg, Germany) and low-retention pipette tips (Corning Inc., Corning, NY, USA) were used to maximize IgG recovery. The samples were filtered via centrifugation (5430 R; Eppendorf, Hamburg, Germany) using centrifuge tube filters with a cellulose acetate membrane and a 0.22 μm pore size (Costar Spin-X; Corning Inc., Corning, NY, USA). Sample concentrations were verified with a UV spectrophotometer (NanoDrop 2000; Thermo Fisher Scientific, Waltham, MA, USA). By measuring the sample absorbance *A* at a wavelength of 280 nm, the antibody concentration *c* in mg/mL was obtained with the Beer-Lambert law, *A* = *∈cl*/*M*, where *∈* = 2.1 ⨉ 10^5^ M^−1^ cm^−1^ is the antibody’s extinction coefficient, *l* = 1 cm is the optical path length, and *M* = 150 kDa is the antibody’s average molecular weight.

### 1.3. Optical Microscopy

The microscopy setup, located within the BioPACIFIC MIP facility in the University of California, Santa Barbara, consisted of an inverted microscope (Axio Observer 7; ZEISS, Oberkochen, Germany) with a computer-controlled and motorized sample stage, a sample incubation system maintained at 25.0 (± 0.3) °C, and a fast digital camera (ORCA-Fire; Hamamatsu Photonics, Hamamatsu, Japan). Samples were loaded into glass capillaries with square cross sections (internal width 500 μm; Friedrich & Dimmock, Millville, NJ, USA). Each capillary was placed on a microscope slide and sealed with epoxy glue at the two ends to prevent solvent evaporation.

For size and *B*_2_measurements, five phase-contrast sequences of 5000 images each were recorded with the ZEISS ZEN 3.9 software and a 20× objective (LD plan neofluar, ZEISS; 0.4 NA, resolution of 0.230 μm/pixel). These image sequences were collected from five distinct (*x*, *y*) positions within the sample, at a focal plane or *z* position roughly in the middle of the capillary to minimize potential boundary effects. A TL halogen lamp light source was used, and videos were acquired at a sampling rate of 500 frames per second, exposure time of 2 ms, image size of 2048(w) ⨉ 256(h) pixels, and depth of focus of 6.88 μm. The imaging conditions and camera settings were maintained throughout this set of experiments. For aggregation stability assessment, size measurements were performed at various times over a span of about 3-4 weeks, with the samples stored at 25.0 (± 0.3) °C.

For viscosity measurements via DDM microrheology, 10 μL aliquots of samples were mixed with fluorescent probe particles (polystyrene particles with a streptavidin-coated surface, yellow, diameter 0.50 μm; Spherotech, Lake Forest, IL, USA) for a concentration of approximately 1.5 ⨉ 10^6^ particles/μL or 0.01 vol%. Three sequences of 1000 images each at three different sample positions were recorded with the ZEISS ZEN 3.9 software and a 20× objective (plan-apochromat, ZEISS; 0.8 NA, resolution of 0.230 μm/pixel). All samples were imaged via epifluorescence to maximize the signal-to-noise ratio, using a Colibri 7 light source and a standard EGFP filter set (excitation at 450-490 nm; emission at 500-550 nm). A frame rate of 10 frames per second was used, exposure time of 100 ms, image size of 512(w) ⨉ 512(h) pixels, and depth of focus of 1.59 μm. All imaging conditions were also kept constant in this set of experiments. Sample measurements exhibiting significant probe aggregation were discarded in further analysis. We also note that some of the microrheology measurements involve videos taken near the side walls of the sample chamber. Although near-wall hydrodynamic interactions are known to affect probe mobility within a distance of ∼10 probe diameters from a solid surface (37), we neglected these effects in the present analysis due to the much larger field of view (>200 probe diameters).

### 1.4. Differential Dynamic Microscopy (DDM)

The experimental workflow is illustrated in **Figure 1**. DDM was performed on the recorded image series using the fastDDM code package, as previously described(22). Briefly, the images are Fourier-transformed, then intensity differences between Fourier-transformed image pairs separated by lag time Δ*t* are computed and ensemble-averaged over the scattering wavevector 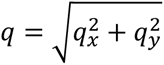 given isotropic structure and dynamics in the sample system. This yields the image structure function *D*(*q*, Δ*t*),

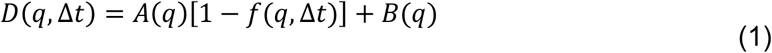

where *A*(*q*) is the static amplitude which represents a convolution of the particle scattering properties and the optical transfer function of the imaging optics, *B*(*q*) is related to the imaging noise and incoherent scattering (which is typically *q* -independent), and *f*(*q*, Δ*t*) is the intermediate scattering function which is related to the same quantity measured by dynamic light scattering (DLS) by an instrumental proportionality constant. As in previous work, *B*(*q*) and *A*(*q*) were estimated from *D*(*q*, Δ*t*) at short and long Δ*t*, respectively (35). For samples of relatively monodisperse particles undergoing Brownian motion, the resulting *f*(*q*, Δ*t*) is expected to be described by a single exponential decay,

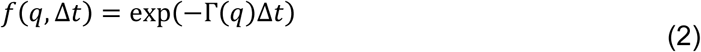

where Γ(*q*) is the characteristic relaxation rate of the sample dynamics at a given *q*.

We emphasize that a single measurement in DDM gives access to a range of *q* values, whereas multiple measurements are required in dynamic light scattering (DLS) to access multiple *q* values which involve physically changing the angle of detection *θθ* for each *q* = (4*πn*/*λ*) sin(*θ*/2), where *n* is the solvent refractive index and *λ* is the wavelength of incident light. In addition, DDM probes much smaller *q* values at ∼0.1-10 μm^−1^, compared to DLS at 22 μm^−1^ (given *θ* = 90° and *λ* = 532 nm), enabling DDM to capture the dynamics of rapidly diffusing particles with reasonable sampling frequencies.

### 1.5. Zeta Potential Measurement

700 μL sample aliquots at 3.8-4.5 mg/mL IgG were filled into folded capillary cells (DTS1070; Malvern Panalytical, Malvern, UK) and their zeta potentials were measured using a Zetasizer (Nano ZS; Malvern Panalytical, Malvern, UK). Three measurements with 10-100 runs each were performed for each sample at a temperature of 25.0 (± 0.1) °C.

### 1.6. Dynamic Light Scattering (DLS)

100 μL sample aliquots at 4.5-6.3 mg/mL IgG were filled into NMR tubes (5 mm outer diameter; Norell, Morganton, NC) and characterized by dynamic light scattering. The instrument (BI-200SM; Brookhaven Instruments, Nashua, NH, USA) is equipped with a goniometer system, a TurboCorr correlator, and a diode-pumped laser at 532 nm (Cobolt Samba, HÜBNER Photonics, Kassel, Germany). The experiments were performed at a scattering angle of 90° and temperature of 25.0 (± 0.1) °C.

## 2. RESULTS

### 2.1. Validating phase-contrast DDM to evaluate protein properties and dynamics

Video acquisition and DDM analysis were performed following the workflow illustrated in **Figure 1**. Phase-contrast imaging was used to perform probe-free DDM, allowing direct characterization of protein-protein interactions, protein sizes, and aggregation without added tracers. Sample viscosities were measured via DDM-based passive microrheology using epifluorescence imaging with fluorescent colloidal probe particles added to the solutions. We first examine probe-free DDM, with **Figure 1** showing a representative frame in a sample video prior to processing. The uneven brightness is attributed to stray light from various sources, such as imperfections on optical surfaces, debris on the optical path, and the surface of the sample cell. Their corresponding contributions to the scattering signal are static or time-invariant, and are therefore removed in the DDM analysis when subtracting the intensities between successive frames separated by lag time Δ*t* (20). Inspection of the image difference in **Figure 1** shows that image subtraction effectively removes the time-independent signal present in the original frames, isolating the signal associated with molecular motion of the antibodies in solution. Since the antibody sizes are far below the diffraction limit, the average speckle size corresponds to the image resolution, while the temporal fluctuations in speckle intensity provide information on protein dynamics and concentration fluctuations across pixels. As Δ*t* increases, more of the objects’ motion is captured leading to a corresponding increase in signal amplitude.

From the image difference (**Figure 1**), the image structure function *D*(*q*, Δ*t*) was calculated, from which the static amplitude *A*(*q*) and the noise *B*(*q*) (Eq. 1) were obtained. Representative plots of these quantities are shown in **Figure 2A** as functions of the scattering wavevector *q*, for 37.1 mg/mL IgG at pH 6. It was observed that *B* is independent of *q* as well as the IgG concentration and buffer condition (**Figure S1**). This is expected, as *B* is related to the imaging noise (e.g., shot noise from the detector) which should be insensitive to the sample details as long as the imaging settings and conditions are kept the same. It was also observed that *A*(*q*) is 1-3 orders of magnitude smaller than *B* depending on the *q* value, demonstrating the challenge in characterizing IgG molecules with hydrodynamic radius *R*_h_ of ∼5-6 nm,(6) which is well below the diffraction limit. This is better understood by examining the resolution function determined by the signal-to-noise ratio *A*_max_(*q*)/*σ*_*B*_, defined here as the ratio between the maximum value of *A* and the standard error *σ*_*B*_ in *B*, shown in **Figure 2B**. We compared the results from phase-contrast imaging used in this work against those from bright-field which was used previously to study various proteins(34). As reflected by the relative magnitudes of *A*_max_(*q*)/*σ*_*B*_, phase-contrast imaging provides a roughly six-fold increase in the signal-to-noise ratio relative to bright-field imaging. This is attributed to the stronger background due to the transmitted light in bright-field imaging (**Figure 2A**), which dominates the detected signal. By comparison, phase-contrast imaging shifts the transmitted light in phase relative to the scattered light and reduces its amplitude, effectively leading to a larger *A*_max_(*q*)/*σ*_*B*_. This difference is clearly visible when comparing normalized images (**Figure 2C–D**) and the corresponding intensity histograms obtained from the two methods (**Figure 2E–F**), where phase contrast imaging exhibits a broader distribution of pixel-wise intensity fluctuations, indicating greater sensitivity to protein concentration fluctuations.

**Figure 2.**
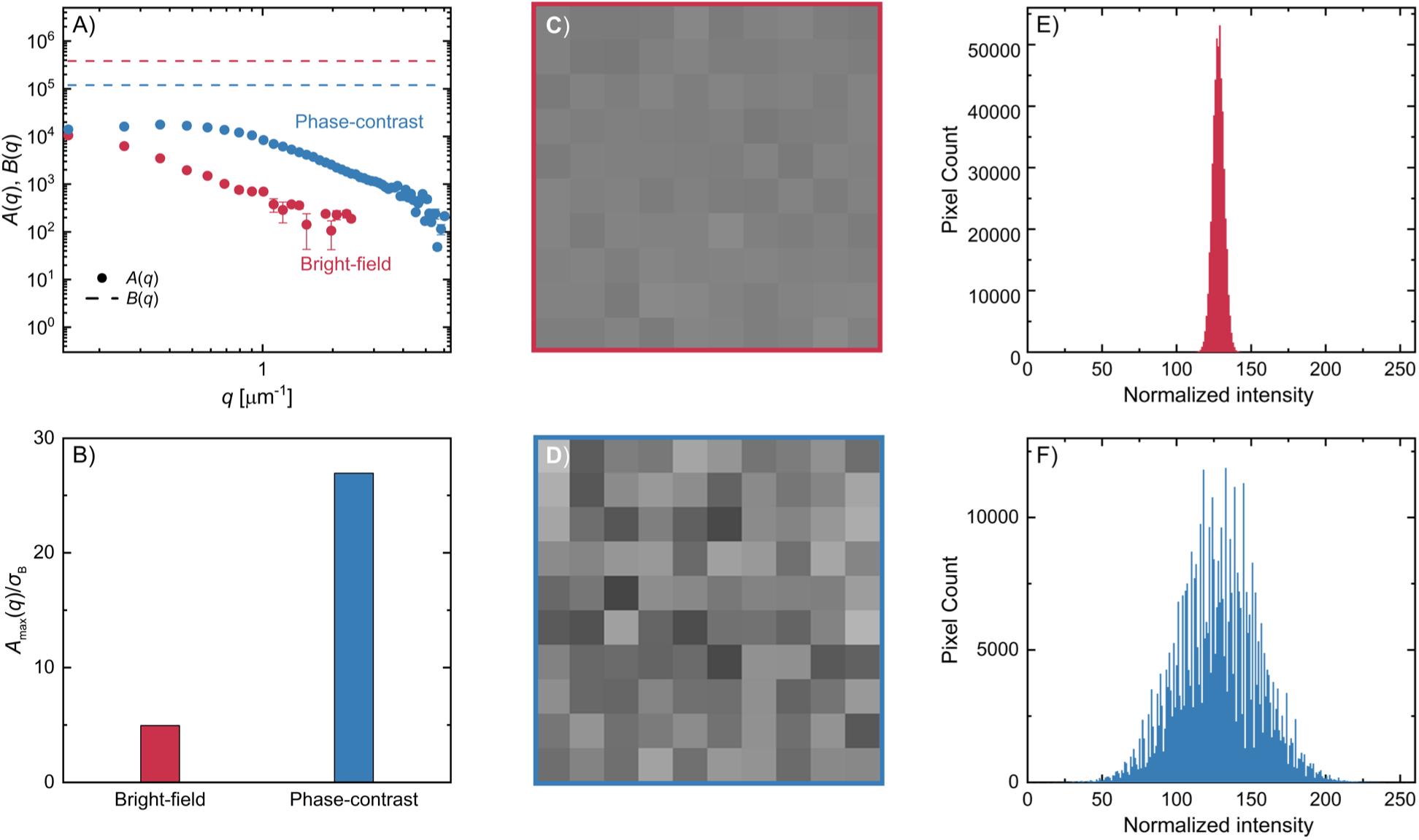
(A) Representative signal amplitude *A*(*q*) (circles) and noise *B*(*q*) (dashed lines) as a function of the scattering wavevector *q* for 37.1 mg/mL IgG in MES buffer at pH 6, shown for bright-field (red) and phase-contrast (blue) imaging modes. (B) Corresponding signal-to-noise ratio *A*_max_(*q*)/*σ*_*B*_, where *A*_max_(*q*) is the maximum value of *A*(*q*) across *q* and *σ*_*B*_ is the standard error of the noise. Error bars represent standard errors from the weighted fit to Eq.. 1. (C) – (D) Image frames from normalized videos (see SI for full videos) of 100 mg/mL polyclonal IgG at pH 6 with 200 mM NaCl in bright-field and phase-contrast microscopy, respectively. A 10 × 10 pixel region of interest (ROI) was selected at the center coordinates of the original videos and the pixel intensities within the ROI were normalized to an identical mean on an 8-bit grey scale. The images have been scaled 25-fold to a display size of 250 × 250 pixels. (E) – (F) Histograms corresponding to the normalized videos for the image frames shown in (C) and (D).

Additionally, the *q*-dependent decay of *A*(*q*) is shifted to higher *q*-values for the case of phase contrast relative to bright-field imaging (**Figure 2A**), improving the resolution of the low-*q* behavior involved in the parameter estimation to follow. Although the current origin of this shift is unknown, it is likely due to the different nature of light-matter interactions encoded in the image intensity between the two imaging modes, and thus a quantitative study of this effect will require theoretical modeling beyond the scope of the present work. To the best of our knowledge, this is the first application of phase-contrast DDM to particles smaller than the diffraction limit, although it has previously been applied to study larger particles such as motile micron-sized microorganisms(32, 38) and anisotropic colloids at hundreds of nm in size(39).

The larger *A*_max_(*q*)/*σ*_*B*_ in phase-contrast imaging effectively decreases the minimum sample concentration that can be successfully analyzed by DDM, and potentially eliminates the need for image post-processing such as 39 to enhance the signal or denoising to artificially increase *A*_max_(*q*)/*σ*_*B*_ (40). The resulting image structure function *D*(*q*, Δ*t*) becomes more well-defined (**Figure S2**), thereby reducing the error associated with downstream data analyses. At smaller *A*_max_(*q*)/*σ*_*B*_, such as that from bright-field imaging in **Figure 2B**, DDM fails to resolve the dynamics in the sample video. This is consistent with a previous study’s empirical signal-to-noise ratio threshold,(34) below which DDM is expected to fail. On the other hand, the upper limit is determined by the onset of multiple scattering from highly turbid samples,(20) which wasn’t a concern in this work since the samples remained transparent up to the highest concentration measured (82.2 mg/mL IgG).

Even in noisy videos (**Figure 1**), where the sample signal is significantly smaller than the image noise (**Figure 2A**) and the sample structures are indiscernible and smaller than the diffraction limit, DDM successfully characterizes the dynamics of the sample through the intermediate scattering function *f*(*q*, Δ*t*) (**Figure 3A**). *f*(*q*, Δ*t*) is well described by a single exponential decay (Eq. 2), indicating a relatively narrow particle size distribution and giving access to the average relaxation rate, Γ(*q*). We emphasize that, in a single measurement, DDM extracts the sample scattering intensity via *A*(*q*), the measurement noise *B*(*q*), and the sample dynamics via Γ(*q*). The *q* -dependence of Γ is shown in **Figure 3B**, in which the observed scaling of Γ ∼ *q*^2^ is consistent with translational Brownian diffusion. By fitting the data with an expression for Brownian diffusion,

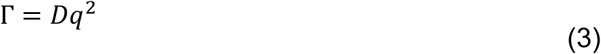

**Figure 3.**
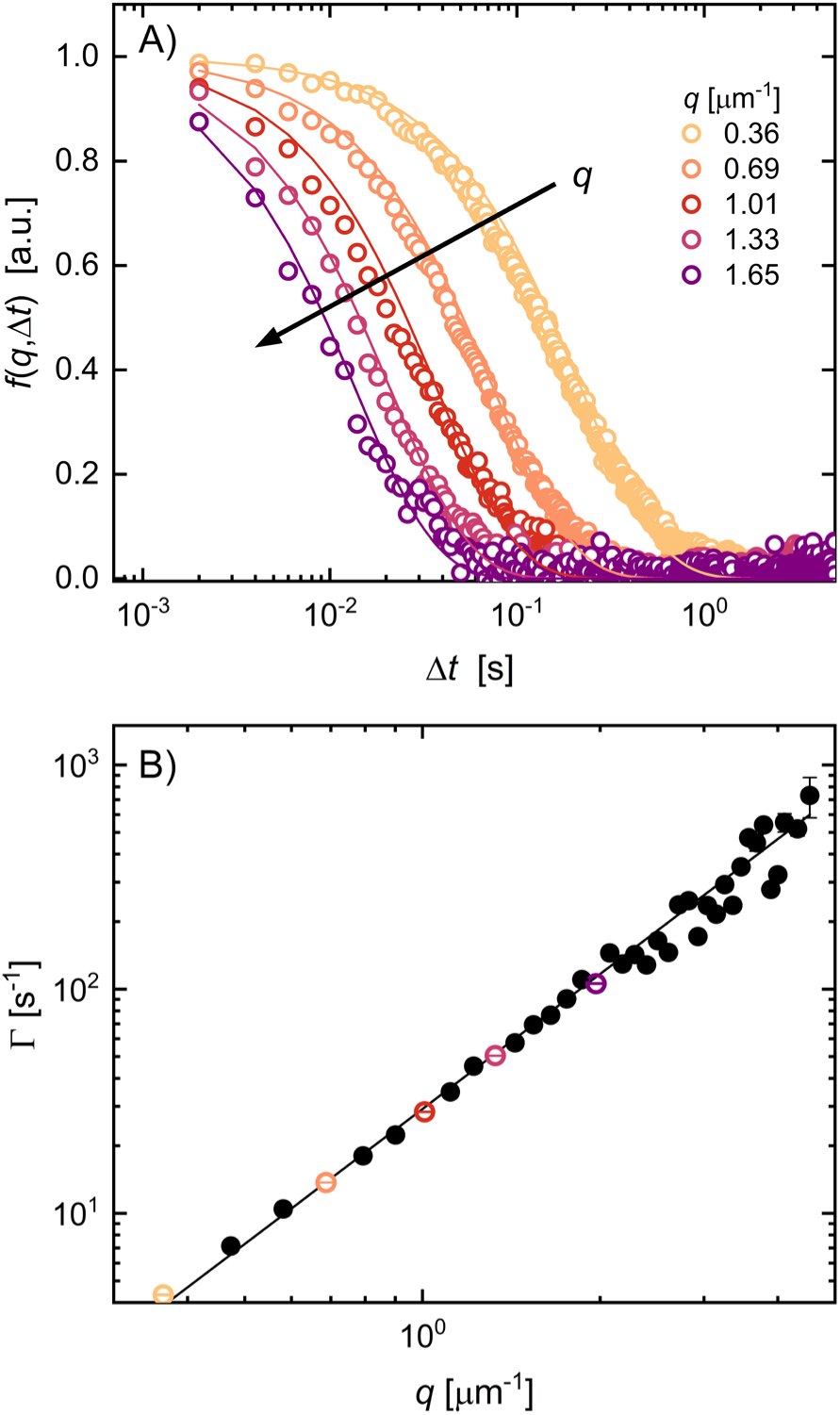
(A) Representative intermediate scattering functions *f*(*q*, Δ*t*) obtained from phase-contrast DDM for 33.3 mg/mL IgG in MES buffer at pH 6 with 100 mM NaCl, shown for five different scattering wavevectors *q*. The experimental data (open circles) are fitted to a single-exponential decay based on Eq. 2 (solid lines). (B) Relaxation rate Γ(*q*) extracted from the fit. Open colored circles highlight the wavevectors *q* corresponding to the curves shown in (A). All remaining Γ(*q*) values are shown as solid black circles. The solid line is a weighted fit to Eq. 3, from which the diffusion coefficient *D* is determined. Error bars represent standard errors from the fits to Eq. 2 (some are smaller than the data markers).

the sample’s concentration-dependent diffusion coefficient *D* was determined. From *D*, the protein-protein interactions, protein size, and solution viscosity via microrheology can be evaluated, as discussed in the subsequent sections.

### 2.2. Characterizing thermodynamic and hydrodynamic protein-protein interactions using DDM

To characterize the thermodynamic protein-protein interactions through the second virial coefficient *B*_2_, the dependence of *A* on *q* and sample concentration *c* was investigated. We refer to the Zimm equation,(41) which expresses the expected asymptotic scattering of particles much smaller than the wavelength of incident light, in the limit of sufficiently small *c* and *qR*_*g*_ ≪ 1, where *R*_*g*_ is the radius of gyration of the protein,

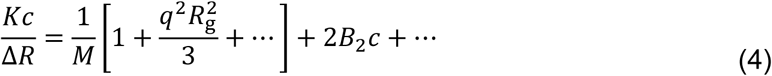

where the higher order terms in *q* and *c* (represented by “…”) are assumed to be negligible. Here, *K* is an optical constant accounting for the refractive index of the solvent and the refractive index increment d*n*/d*c* of the protein in solution, Δ*R* is the Rayleigh ratio which encodes the measured scattering intensity, *M* is the protein’s molecular weight. As in the work of Guidolin *et al.*,(34) to apply Eq. 4 to DDM analysis, we assume for any given *q*-value that Δ*R* is directly proportional to the static intensity amplitude factor *A*, i.e., Δ*R*(*q*) ∝ *A*(*q*). With this assumption, and in the limit of *q* → 0, Eq. 4 reduces to

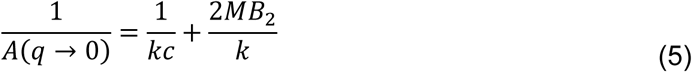

where *k* is a new constant that accounts for the optical transfer function of the microscope, in addition to *K* and the linear proportionality constant between Δ*R* and *A*(*q*) (20). The extrapolated value *A*(*q* → 0) is determined by performing a linear regression of *A*(*q*) corresponding to **Eq. 5** in the *q*-range spanning 0.4 − 1.0 *μ*m^−1^. In cases where low-*q* data contained clear outliers (1-2 orders of magnitude larger than the surrounding values), the lower bound of the fitting range was adjusted accordingly to exclude those points.

**Figure 4** shows a representative linearized plot of 1/*A*(*q* → 0) vs 1/*c* (results for other sample conditions in **Figure S3**). Fitting the data to Eq. 5 using an average value of *M* = 150 kDa provides an estimate for *B*_2_ for a given set of formulation conditions. The resulting *B*_2_ values are shown in **Figure 5A** for different pH with 100 mM NaCl, and in **Figure 5B** for different concentrations of added NaCl at pH 6. We first note that the positive *B*_2_ values in **Figure 5** reflect net repulsion-dominated protein-protein interactions, whereas negative values would reflect attraction-dominated interactions (42). These *B*_2_ values from DDM agree well with previously reported values of ∼4×10^−5^ to 3×10^−4^ mol mL g^−2^ for monoclonal IgG1, IgG2, and IgG4 under similar sample conditions using either static light scattering(43, 44) or a combination of DLS and analytical ultracentrifugation (11). The human polyclonal IgG used in this work is comprised of four IgG subclasses, IgG1 to IgG4, with IgG1 accounting for ∼60-70% relative abundance in adult human serum (45). Therefore, the values reported here should be interpreted as averages across the binary interactions among these subclasses.

**Figure 4.**
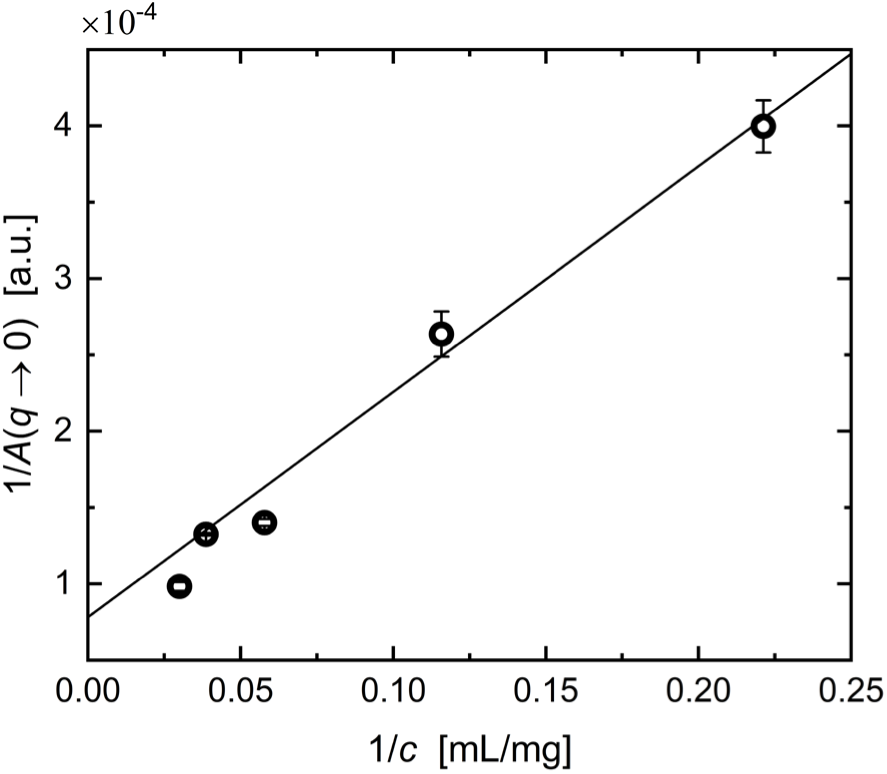
Representative linearized plot of DDM amplitude *A*(*q* → 0) (circles) versus protein concentration for IgG in MES buffer at pH 6 with 100 mM NaCl. Error bars reflect the standard errors from the linear regression used to extrapolate *A*(*q*) to *q* → 0. The solid line represents a weighted fit based on Eq. 5.

**Figure 5.**
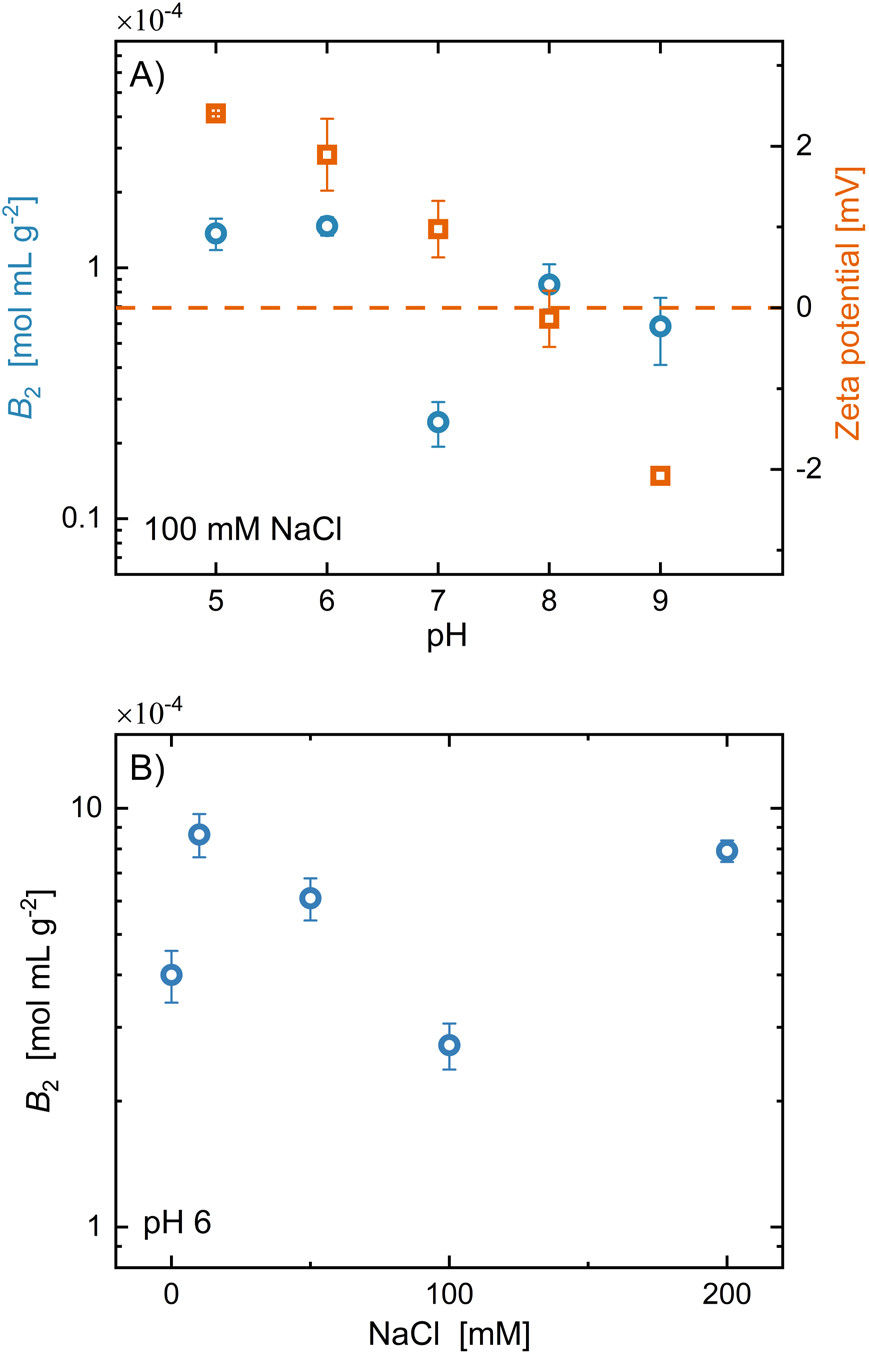
Second virial coefficient *B*_2_ (blue circles) as a function of (A) sample pH with 100 mM NaCl and (B) NaCl concentration at pH 6 in 20 mM histidine buffer. Different buffer systems were used in (A) and (B) to reflect experimental design; buffers in (A) were selected to match the target pH. Orange squares in (A) represent the zeta potential, with the dashed line indicating the point of zero charge (PZC), where the zeta potential is zero and the net surface charge on IgG is neutral. Error bars on *B*_2_reflect standard errors from the weighted fits to Eq. 5, while those on the zeta potential indicate the standard error of the mean from three replicate measurements.

A general non-monotonic dependence of *B*_2_ on both sample pH (**Figure 5A**) and salt concentration (**Figure 5B**) was observed. For the case of pH-dependent measurements, the minimum in *B*_2_at pH 7 roughly coincides with the measured point of zero charge (PZC), corresponding to a net zero surface potential on the antibodies as indicated by zeta potential measurements (right axis on **Figure 5A**; additional measurements at different pH without added salt can be found in **Figure S4**). The zeta potential values here agree well with reported values^12,45^. The pH-dependence of *B*_2_ also correlates with the related isoelectric point (pI), corresponding to net zero overall charge on the antibodies, which was reported to be approximately 7-9 in a previous study,(47) measured by capillary isoelectric focusing and also calculated from primary sequences for IgG1. The wide range in this reported pI value is due to heterogeneity in the antigen-binding site of IgG1 in the mixture, which arises from different primary structures and thus varied pI values within the same IgG1 subclass (47). Nonetheless, we infer from these results that the measured trends in *B*_2_can be explained by expected differences in screened electrostatic repulsions arising from changes in pH and ionic strength of the protein solutions. As the pH approaches the PZC, the net charge on the surface of the antibody particles decreases, resulting in weaker protein-protein repulsions and thus lower *B*_2_ values (**Figure 5A)**.

A similar trend was observed in *B*_2_ with pH in the absence of added salt (**Figure S4**), and in related studies (4, 44). The observed decrease in *B*_2_ with increasing NaCl concentration (**Figure 5B**) is consistent with increased screening of electrostatic repulsions with increasing ionic strength of the solvent. The minimum in *B*_2_ at 100 mM NaCl is also consistent with previous findings,(11, 43) although the subsequent increase at 200 mM NaCl is inconsistent with theories for screened electrostatic interactions of globular colloids. This observation likely reflects the challenges typically associated with measuring screened electrostatic interactions arising from ion-ion correlations at relatively high salt concentrations (48).

Using the same dataset as the *B*_2_measurements, the diffusion interaction parameter *k*_d_ was determined from the concentration-dependence of the diffusion coefficient *D* (representative data shown in **Figure 6**; results for other sample conditions in **Figure S5**). *k*_d_ is a more extensively studied measure of protein-protein interactions, and encodes both thermodynamic and hydrodynamic interactions between proteins. The DDM results agree well with the measured value of *D* = 34.1 μm^2^ s^−1^ from DLS for 32.6 mg/mL IgG under the same solution conditions. In the limit of sufficiently weak protein-protein interactions, the dependence of *D* on *c* can be described by(49, 50)

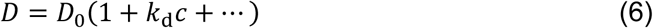

**Figure 6.**
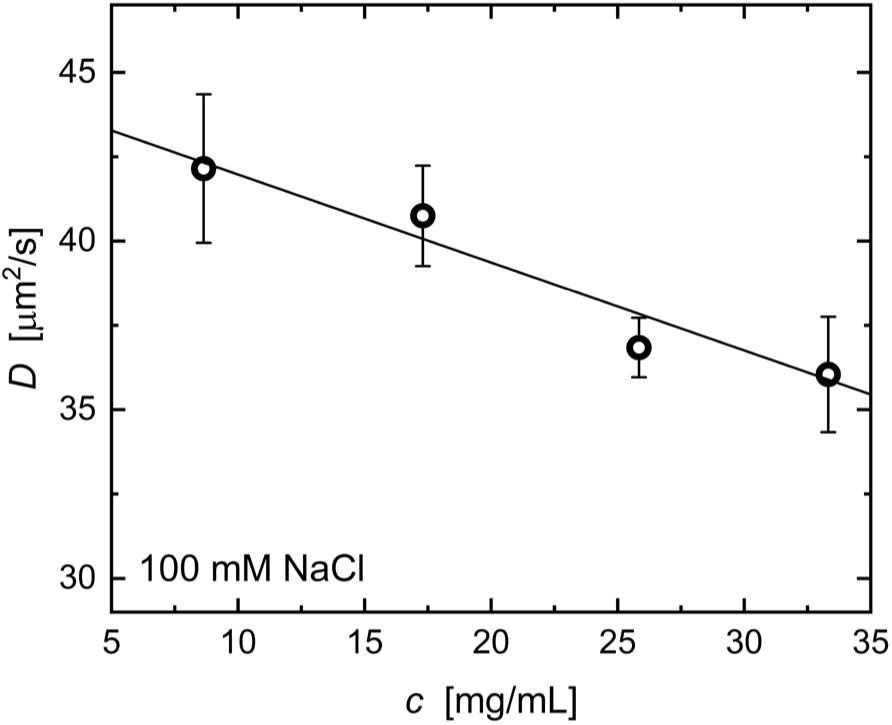
Representative diffusion coefficient *D* (circles) as a function of IgG concentration *c* in MES buffer at pH 6 with 100 mM NaCl. Error bars represent standard errors determined from the weighted fit to Eq. 3. The solid line shows a weighted fit based on Eq. 6.

where *D*_0_ is the self-diffusion coefficient at infinite dilution. Note that the higher order terms in the virial expansion in *c* in Eq. 6 are assumed to be negligible under the relatively low dilute conditions studied here. The resulting *k*_d_ values are shown at different pH with 100 mM NaCl in **Figure 7**. *k*_d_ at different salt concentrations could not be accurately determined because of the negligibly small dependence of *D* on protein concentration under these conditions. The *k*_d_ values in this work are within the wide range of reported values in the literature from −40 to 70 mL g^−1^ for monoclonal IgG1, IgG2, and IgG4 under similar sample conditions using DLS (7, 13).

**Figure 7.**
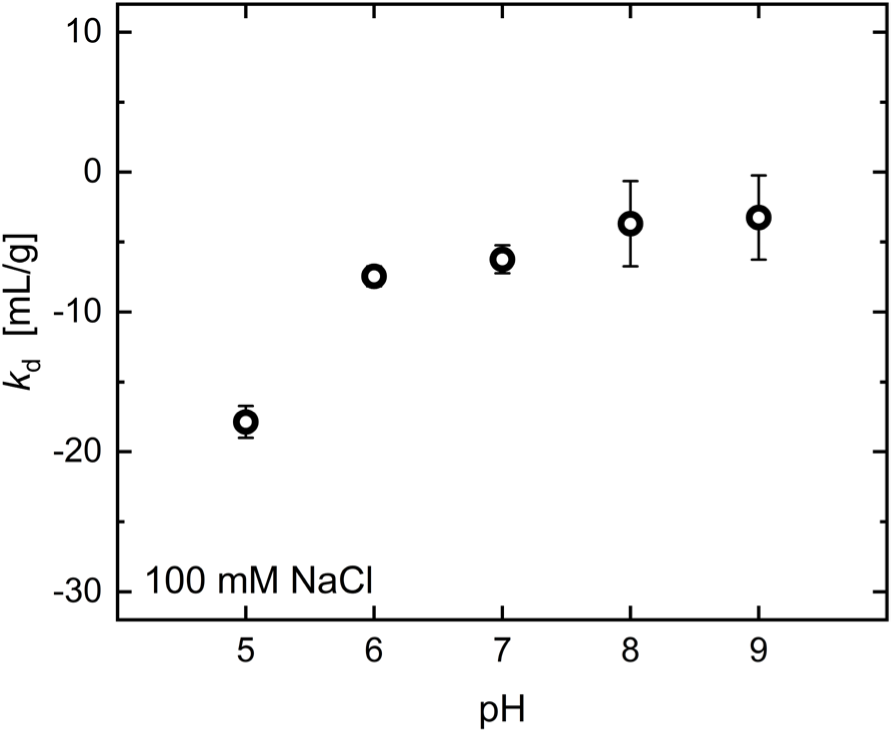
Diffusion interaction parameter *k*_d_ as a function of sample pH with added 100 mM NaCl. Error bars represent standard errors from the weighted fit to Eq. 6.

In the present work, we observed negative values of *k*_d_ over the entire range of measured conditions. This same observation of *B*_2_ > 0 and *k*_d_ < 0 has been made previously for monoclonal IgG1 and IgG2 (4, 11). It is important to emphasize that *k*_d_ incorporates non-dilute effects of protein-protein interactions on diffusive dynamics arising separately from thermodynamic interactions encoded by *B*_2_, hydrodynamic interactions due to solvent flow between diffusing molecules, and entropic interactions due to the excluded volume of nearby protein molecules. In the limit of low concentrations, these three contributions are linearly separable, leading to(49)

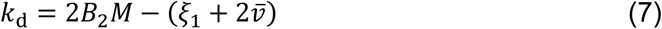

where *ξ*_1_ represents the leading-order contribution to the friction coefficient of an individual protein molecule due to hydrodynamic interactions, and *v̅* is the specific volume of the protein molecule (representing entropic interactions). Since *ξ*_1_and *v̅* are strictly positive and *B*_2_is positive in this work under all studied conditions, the sign of *k*_d_ thus depends on the relative magnitudes of these parameters in Eq. 7. The negative *k*_d_ values observed in **Figure 7** therefore indicate that hydrodynamic and entropic protein-protein interactions dominate over (enthalpic) thermodynamic interactions for the range of conditions measured. As a result, the trend in *k*_d_ is not expected to correlate well with the observed trend in *B*_2_.

### 2.3. Characterizing protein size, aggregation, and solution viscosity using DDM

On the same set of low-volume sample solutions, DDM can determine both the apparent protein hydrodynamic radius *R*_h_ and the solution viscosity using the Stokes-Einstein-Sutherland equation,

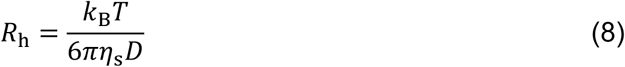

where *k*_B_ is the Boltzmann constant and *T* is the absolute temperature. Using the measured solvent viscosity *η*_*s*_ and diffusion coefficient *D* of the antibody molecules from DDM, *R*_h_ of the antibody particles can be directly estimated from Eq. 8. To assess the aggregation propensity of the protein solutions under formulation-relevant conditions, we measured *R*_h_ values over time on samples at high IgG concentrations of 74.2-82.2 mg/mL stored at 25.0 (± 0.3) °C at different pH with 100 mM NaCl (**Figure 8A**), and different concentrations of added NaCl at pH 6 (**Figure 8B**; corresponding Γ(*q*) data in **Figure S6**). The *R*_h_ values from DDM agree well with those from DLS (indicated by the solid line). *R*_h_ showed no systematic dependence on sample pH and NaCl concentration. No change in *R*_h_ was observed over a span of 3-4 weeks for any of the studied conditions, indicating no measurable aggregation. This suggests that the commercially available polyclonal IgG antibody remained colloidally stable and well dispersed over time without significant aggregation, sedimentation, or adsorption onto the glass capillary walls.

**Figure 8.**
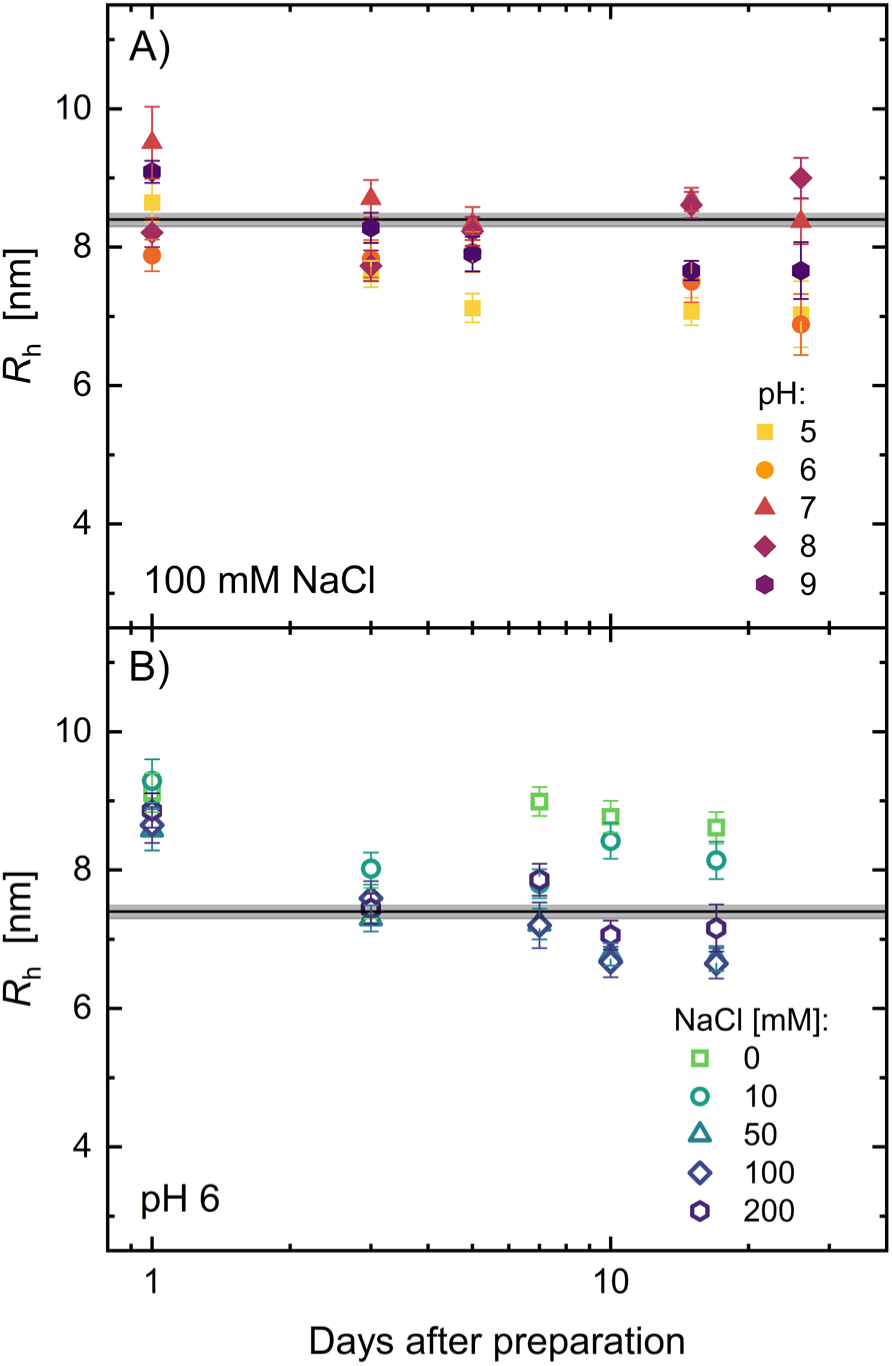
Time evolution of hydrodynamic radius *R*_h_ measured at IgG concentrations ranging from 74.2 to 82.2 mg/mL under (A) varying pH conditions with 100 mM added NaCl (solid symbols) and (B) varying NaCl concentrations at pH 6 in 20 mM histidine buffer (open symbols). Different buffer systems were used in (A) and (B) to align with the experimental design; buffers in (A) were selected to match the target pH. Error bars represent standard errors determined from weighted fits to Eq. 8. In (A), the solid line represents the average *R*_h_ measured by DLS for 6.3 mg/mL IgG at pH 8, while in (B), it corresponds to that for 4.5 mg/mL IgG without added salt. The shaded gray region around the solid line indicates the standard error of the mean from three replicate measurements.

### 2.4. Testing colloidal models for protein solution viscosity

The viscosities of the same samples of the previous sections were measured using passive microrheology experiments coupled with DDM analysis (**Figure 1**), in which fluorescent colloidal probe particles were added to the protein samples and imaged with epifluorescence microscopy (experimental details in Section 1.3). Using the measured value of *D* for the probe particles from DDM (Eq. 3), and the known values of *R*_h_ for the probes provided by the manufacturer, the sample viscosity *η* was determined (Eq. 8) and is shown in **Figure 9** (corresponding Γ(*q*) data in **Figure S7**). DDM-based viscosity measurements were previously benchmarked against the more conventional techniques of bulk rheology and capillary viscometry, producing good quantitative agreement (29). Similar viscosities have been observed for samples of monoclonal IgG at 150 mg/mL depending on the specific antibody type (13). The lack of instabilities in the samples, in the form of aggregation and high sample viscosities, can be correlated with the positive *B*_2_ values (**Figure 5**), which again indicate net repulsive protein-protein interactions resulting in electrostatic stabilization of the antibody molecules. These interactions are evident in the viscosity, which exhibits values significantly above those expected from the general correction to the solvent viscosity for suspensions in the presence of nondilute long-range hydrodynamic interactions(51),

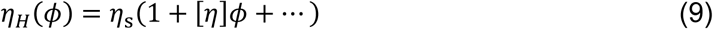

**Figure 9.**
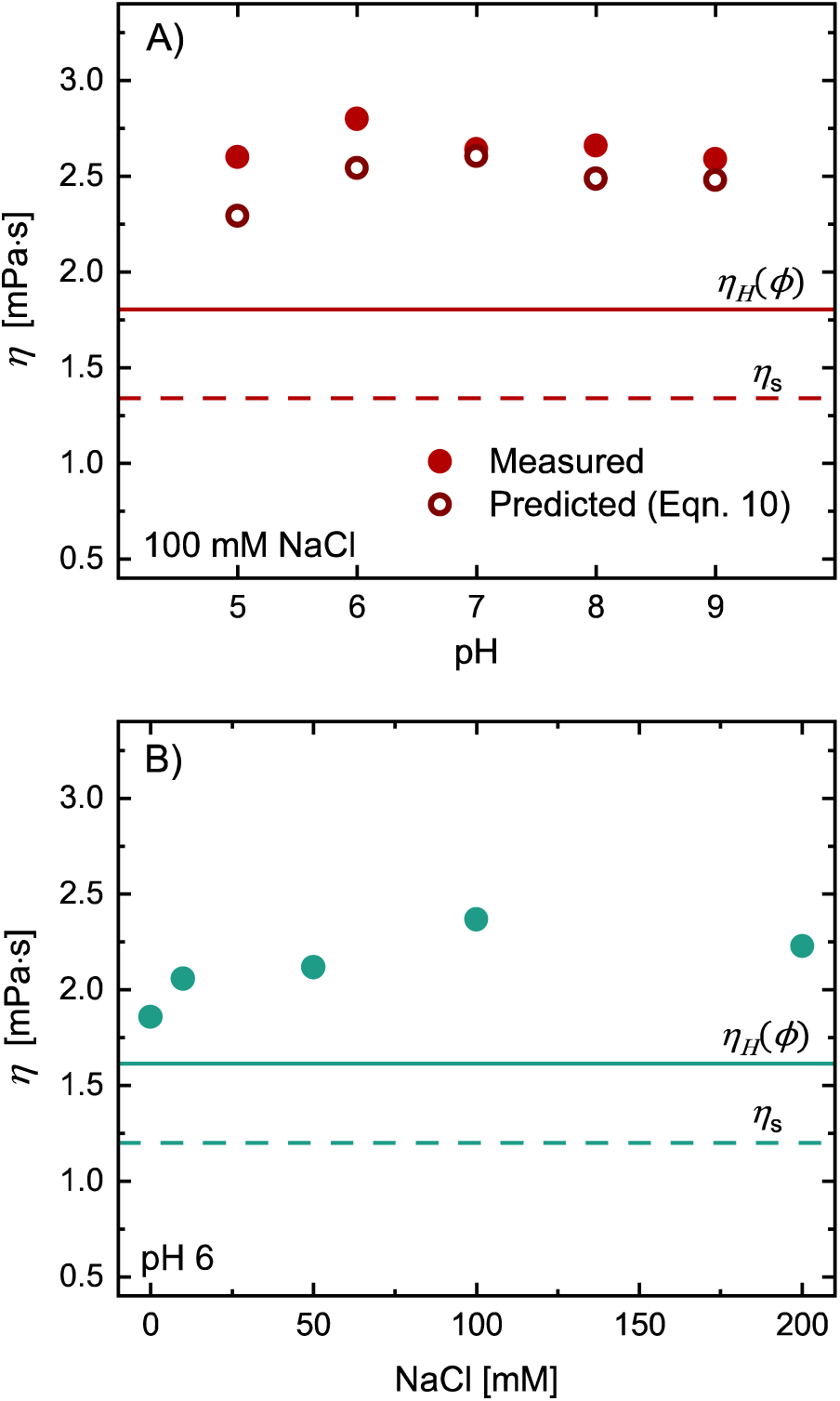
Viscosity *η* of IgG measured at concentrations ranging from 74.2 to 82.2 mg/mL under (A) varying pH with 100 mM added NaCl (solid circles), and (B) varying NaCl concentrations at pH 6 in 20 mM histidine buffer (open circles). In (A), the open red circles correspond to the predicted viscosity based on the colloidal model proposed by Godfrin *et al.* (18) (**Eq. 10**). In both (A) and (B), solid lines indicate the predicted hydrodynamic viscosity contribution *η*_*H*_(*ϕ*) (**Eq. 9**). The dashed red line in (A) corresponds to the viscosity of 25 mM Tris buffer at pH 7 while the dashed green line in (B) shows that of 20 mM histidine buffer at pH 6. Different buffer systems were used in (A) and (B) to align with the experimental design; buffers in (A) were selected to match the target pH. Error bars represent standard errors calculated from weighted fits to **Eq. 8** (some are smaller than the data markers).

where *ϕ* is the protein volume fraction (which we estimate assuming a mass density of 1.353 g/cm^3^) (52) and [*η*] is the intrinsic viscosity. A value of [*η*] = 9 mL/g was used for all calculations, representing an average of values previously reported for different IgG variants (53). The deviation of the measured viscosities (filled symbols in **Figure 9**) from predictions of **Eq. 9** (solid lines in **Figure 9**) result suggests the importance of nondilute thermodynamic interactions or other effects involving antibody shape and deformability in determining the solution viscosity at these relatively high concentrations (*ϕ*∼0.056).

To demonstrate the utility of intensified DDM measurements to characterize protein formulation properties in solution, we use the viscosity measurements to self-consistently test a colloidal theory for the solution viscosity. Godfrin *et al.* (18) recently presented a simplified colloidal model to account for these effects, in which protein-protein interactions are modeled by pairwise adhesive interactions between effective hard spheres. Assuming linear separability of contributions to the solution viscosity arising from hydrodynamic interactions, protein-protein self-attractions (e.g., due to nonspecific associations), and protein-protein self-repulsions (e.g., due to electrostatic interactions), they proposed the following model for the reduced viscosity, *η*_*r*_ = *η*⁄*η*_*s*_,

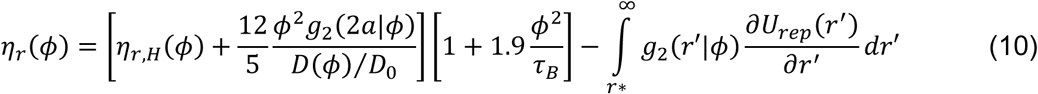

where *η*_*r*,*H*_(*ϕ*) = *η*_*H*_(*ϕ*)⁄*η*_*s*_ is a hydrodynamic contribution due to monomeric proteins, *g*_2_(*r*|*ϕ*) is the concentration-dependent pair radial distribution function (with *g*_2_(2*a*|*ϕ*) being its value at protein-protein contact), *D*(*ϕ*) and *D*_0_are the concentration-dependent and limiting dilute diffusivities as previously defined, *τ*_*B*_ is the so-called Baxter “stickiness” parameter characterizing the strength of protein-protein adhesion, and *U*_*rep*_(*r*) is a pair interaction potential characterizing long-range protein-protein repulsions. The first bracketed term in **Eq. 10** comprises a hydrodynamic contribution which corrects that arising from pure hard sphere interactions with a contribution arising from small transient clusters formed due to protein-protein attractions. The second bracketed term comprises a Brownian contribution that accounts for the hindered mobility of individual protein molecules due to transient adhesive interactions. The integral term accounts for nondilute thermodynamic interactions due to protein-protein repulsions. We will ultimately show that this latter term is unimportant for the conditions we study, and so it will be ignored in the subsequent calculations.

Godfrin *et al.* (18) tested predictions of **Eq. 10** using independent measurements of the various parameters on the right-hand side via neutron scattering methods. We instead show here that – in the case where the effects of protein-protein repulsions can be ignored – all of the relevant parameters in **Eq. 10** can be estimated using the DDM measurements presented in the previous sections. To estimate *η*_*r*,*H*_(*ϕ*), we use **Eq. 9**, i.e., *η*_*r*,*H*_(*ϕ*) = 1 + [*η*]*ϕ*. The quantity *D*(*ϕ*)⁄*D*_0_ is obtained directly from the probe-free DDM measurements in **Section 2.2** via **Eq. 6**. The quantity *τ*_*B*_ for the Baxter adhesive sphere model can be obtained directly from the measured values of the second virial coefficient *B*_2_ estimated from probe-free DDM measurements via(18)

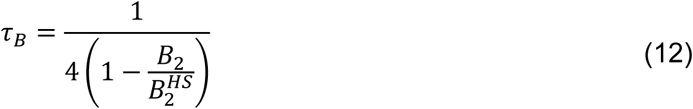

where 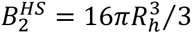 is the virial coefficient for an equivalent hard sphere of radius equal to the measured hydrodynamic radius of a monomer (obtained here from DDM). Finally, we estimate *g*_2_(2*a*|*ϕ*) using the theoretical treatment of Chiew and Glandt(54) for adhesive hard spheres at contact, where

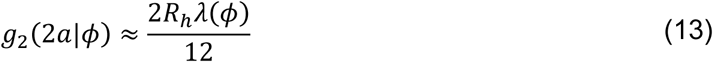

The value of *λ*(*ϕ*) is given by the positive root of the following equation,

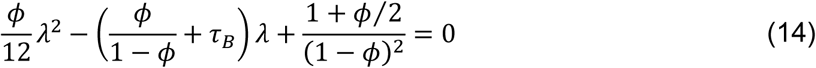

We emphasize that, upon neglecting the effects of protein-protein repulsions, **Eq.s 9-14** comprise a closed set of equations by which to predict the viscosity from the parameters *R*_*h*_, *B*_2_, *k*_d_ and [*η*], and that each one of these four independent parameters can be extracted from the DDM measurements and analysis presented in this work.

**Figure 9A** compares viscosity predictions made using **Eq. 10** (open symbols) to the experimentally obtained values of pH-dependent viscosity for IgG at a concentration of *ϕ*∼0.08 and 100 mM NaCl (closed symbols). Overall, we find a remarkable level of agreement between the predictions and measurements, especially when considering that the model predictions contain no adjustable fitting parameters. This includes the weak non-monotonic pH-dependence of *η*, which is captured by the model due to the observed non-monotonic pH-dependence of *B*_2_(**Figure 5A**). We note that the model calculations underscore the importance of the effects arising from protein-protein attractions, since the observed viscosities are significantly greater than those predicted solely based on the hydrodynamic contribution from individually-dispersed proteins through *η*_*r*,*H*_(*ϕ*) alone (solid lines in **Figure 9**). However, we do note the tendency for the model to systematically underpredict the viscosity by as much as 15%. This finding is consistent with the previous results of Godfrin *et al*(18)., and verifies that the viscosity contribution due to protein-protein repulsions is unimportant, since the integral term in **Eq. 10** is strictly negative and therefore can only serve to decrease the predicted viscosity. Nevertheless, the ability to compare measurements and model predictions for the solution viscosity using parameters obtained entirely from the DDM measurements demonstrates that the intensified workflow developed in this work can be used to self-consistently evaluate molecularly-informed models for the viscosity of protein formulations that capture the effects of thermodynamic and hydrodynamic protein-protein interactions.

## 3. DISCUSSION

We have demonstrated the ability of DDM to successfully characterize the hydrodynamic radius *R*_h_, second virial coefficient *B*_2_, diffusion interaction parameter *k*_d_, and sample viscosity *η* of antibody samples from a concentration series of low-volume samples using simple video microscopy experiments. Good quantitative agreement was observed between the values of these parameters obtained from phase-contrast DDM and the values independently measured either in this work or in previous reports using more conventional techniques. The ability to measure all of these properties from a single series of measurements using DDM allowed us to self-consistently test a predictive theory for the solution viscosity (**Eq. 10**), which showed reasonable agreement with measured viscosities without any adjustable parameters.

We emphasize that the resources consumed by the DDM experiments are significantly less than what would have been required using conventional measurements of the relevant protein and solution properties. To quantify this, comparisons of amount of protein consumed, measurement time, and measured value between DDM and these conventional techniques are shown in **Table I**. When evaluating the full set of parameters (*R*_h_, *B*_2_, *k*_d_, and *η*) on the same sample sets (DDM determines *B*_2_ and *k*_d_ on the same samples and dataset; the same for SLS/DLS), DDM requires only ∼3 mg of sample and ∼2 minutes of total measurement time whereas a combination of three conventional techniques (SLS, DLS, viscometry) required 120 mg of sample and ∼40 minutes of total measurement time. The decrease in both the required sample amount and measurement time by nearly two orders of magnitude using DDM over other techniques highlights the high-throughput capabilities of DDM in screening antibody formulations.

**Table I.** Comparison of amount of protein consumed, measurement time, and measured or previously reported value between differential dynamic microscopy (DDM) and conventional techniques.

| Biophysical property | Measurement technique | Protein consumed (mg) | Measurement time (min) | Measured or reported value |
| --- | --- | --- | --- | --- |
| $B_2$ (mol mL g<sup>-2</sup>)<sup>a</sup> | DDM | 1 | 1 | $5.5 \times 10^{-6}$ to $8.7 \times 10^{-4}$ |
| | SLS | 10 | 30 | - $2.0 \times 10^{-4}$ to $2.0 \times 10^{-4}$ (43) |
| | AUC-SE | 2 | 1440–2880 | - $3.2 \times 10^{-5}$ to $2.1 \times 10^{-5}$ (4) |
| $k_d$ (mL/g)<sup>b</sup> | DDM | 1 | 1 | -18 to -4 |
|  | DLS | 10 | 10 | -40 to 70 (13) |
| $R_h$ (nm)<sup>b</sup> | DDM | 1 | 0.2 | 6.7 to 9.5 |
|  | DLS | 10 | 1 | 7.2 to 8.4 |
| $\eta$ (mPa s)<sup>c</sup> | DDM | 1 | 0.2 | 1.8 to 2.8 |
|  | Viscometry | 100 | 5 | ~1 to 250 (13) |

<sup>a</sup>Second virial coefficient $B_2$ is conventionally measured using static light scattering (SLS), or analytical ultracentrifugation–sedimentation equilibrium (AUC–SE).
<sup>b</sup>Diffusion interaction parameter $k_d$ and the hydrodynamic radius $R_h$ are conventionally measured with dynamic light scattering (DLS).
<sup>c</sup>Sample viscosity is conventionally measured with viscometry.
*Note:* For $B_2$ and $k_d$ measurements, five protein concentrations (5–40 mg/mL) were considered to estimate the sample amount and measurement time for all the listed techniques, except for AUC–SE where only one concentration (10 mg/mL) was considered, in accordance with a typical experiment(4). For $R_h$ and $\eta$ measurements, only one protein concentration (100 mg/mL) was considered for all techniques.

By combining multiple characterization workflows into a single measurement, DDM enables the simultaneous interpretation of molecular and macroscopic-scale properties. When applied to the polyclonal antibody solutions studied here, DDM reveals that the pH-dependent values of *B*_2_ exhibited non-monotonic behavior with a minimum that agreed well with the measured point of zero charge for the antibodies, and a dependence on monovalent salt concentration that reflected the expected screening of electrostatic repulsions. Nevertheless, the estimated *B*_2_ values always remain positive, indicating net repulsive protein-protein interactions for a wide range of solution pH and ionic strength, suggesting successful electrostatic stabilization of the protein. By contrast, the negative measured values of *k*_d_ suggest that hydrodynamic interactions are significant for the range of protein concentrations studied. Overall, these findings suggest that the formulations studied here should remain stable against aggregation and high viscosities, and this is confirmed by the measurements of *R*_h_ and *η* for the protein solutions. The utility of DDM measurements to characterize protein formulation properties in solution goes beyond qualitative interpretations, as demonstrated by the ability to use DDM to self-consistently test molecular-level theory for the viscosity of protein formulations. We find remarkable agreement between the viscosities measured with DDM and those predicted by the colloidal model developed by Guidolin *et al*., which captures the effects of thermodynamic and hydrodynamic protein-protein interactions.

Although the experiments here were conducted on a well-formulated commercial antibody with relatively high stability, we anticipate that the same approach can be applied to less-optimized formulations. Notably, the signal-to-noise ratio in phase contrast DDM is expected to increase for proteins with stronger interactions or higher aggregation propensity, since these conditions will produce larger fluctuations in image intensity, thereby improving measurement precision (34).

## 4. CONCLUSIONS

This work established the potential for optical microscopy in concert with DDM analysis to intensify conventional biophysical measurements of protein solutions, including measures of protein-protein interactions (both thermodynamic and hydrodynamic), protein size, aggregation stability, and solution viscosity, which are critical for formulation development. The accuracy of DDM to evaluate these properties was assessed using experiments on polyclonal antibody formulations over a wide range of conditions. Phase-contrast imaging was used to collect videos of the antibody samples, providing nearly an order of magnitude increase in the signal-to-noise ratio relative to conventional bright-field imaging. This effectively decreases the lowest measurable concentration and eliminates the need for post-processing of videos to artificially improve the measurement precision(40).

With smaller sample requirements and shorter measurement times, DDM offers higher throughput access to *B*_2_ and *k*_*D*_ than is typically achieved with conventional techniques. Additionally, DDM also enables testing molecular-level models that capture thermodynamic and hydrodynamic interactions of proteins, providing a self-contained method for connecting protein-protein interactions to measured formulation properties. Its simultaneous access to other industrially relevant parameters, such as particle size and aggregation, as well as solution viscosity, provides a roadmap to correlate *B*_2_ and *k*_*D*_ with these more readily observable indications of formulation instability, and potentially allow their prediction early during formulation development, all within the same experimental workflow. Combined with its potential for full automation (i.e. using autosamplers and pre-programmed video collection) and iterative screening protocols (i.e. using machine learning to inform the next set of conditions that would lead to the target formulation properties),(30) we expect that DDM is poised to become an attractive routine technique to rapidly develop protein formulations for biopharmaceutical applications.

## DATA AVAILABILITY

Datasets for this work, including numerical data for all figures and representative raw video files are available through the Dryad data repository, [DOI to be entered here upon publication of the work].

## AUTHOR CONTRIBUTIONS

C.I.G., A.S., M.T.V., and M.E.H. conceived the study and developed the methodology. C.I.G. and A.S. carried out the experiments and performed data curation, analysis, and visualization. J.M.U. contributed to data analysis and software development. M.T.V. and M.E.H. supervised the project and provided resources. N.M. and R.G. provided conceptual guidance, project administration, and funding. C.I.G. and A.S. wrote the original draft. M.T.V., and M.E.H. edited the manuscript.

## ACKNOWLEDGMENTS

We acknowledge generous support from Hamamatsu and Carl Zeiss Microscopy for the use and installation of the demonstration camera used in this work, and thank John Parsons, Mark Mobilia, Mark Fairchilds, and Geert Vreede for technical assistance. We also acknowledge Enrico Lattuada and Roberto Cerbino for the development and helpful discussions regarding use of the fastDDM software package. This work was primarily funded by the BASF California Research Alliance (CARA, UCB 052673/SA-10916), with partial support from the MRSEC Program of the National Science Foundation (NSF) under Award No. DMR 2308708 (IRG-2). This work made use of the NSF-supported BioPACIFIC Materials Innovation Platform (DMR-2445868) and Biological Nanostructures Laboratory within the California NanoSystems Institute, supported by the University of California, Santa Barbara and the University of California, Office of the President.

## SUPPLEMENTAL INFORMATION

Supplementary information is available online and includes seven supplemental figures (Figures S1–S7) and two representative microscopy videos.

**“SI_02_BrightField” and “SI_03_PhaseContrast”** show representative videos of 100 mg/mL polyclonal IgG at pH 6 with 200 mM NaCl, recorded using bright-field and phase-contrast microscopy, respectively. A 10 × 10 pixel region of interest (ROI) was selected at the center coordinates of the 2048(w) × 512(h) pixel original videos and the pixel intensities within the ROI were normalized to an identical mean on an 8-bit grey scale. For improved visualization, the videos were scaled 25-fold to a display size of 250 × 250 pixels. Each video was truncated to half of its original number of frames. These videos correspond to the histograms in **Figure 2E– F.**

