## Supplemental Information for "Measurement intensification for antibody formulations: combined measurement of protein size, interactions, and viscosity by differential dynamic microscopy"

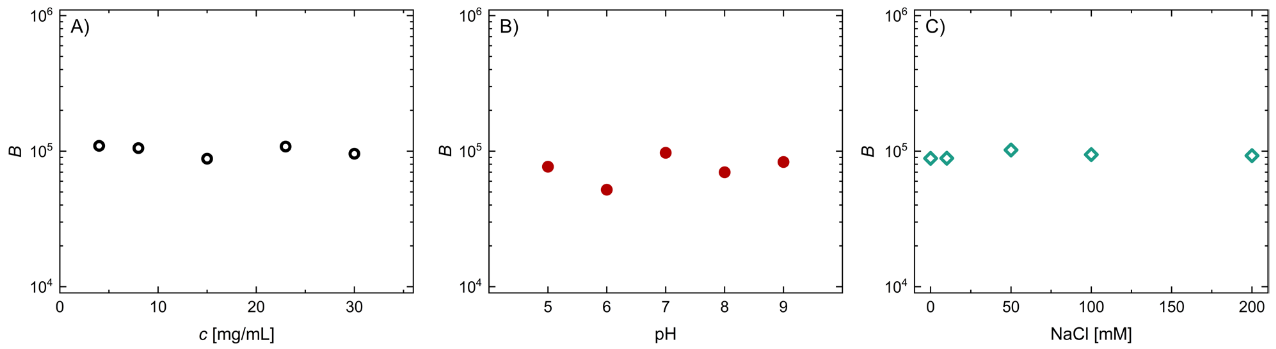


**Figure S1**. Noise $B$ as a function of (A) varying IgG concentrations $c$ in MES buffer at pH 6 with 100 mM NaCl, (B) varying pH conditions with 100 mM added NaCl, and (C) varying NaCl concentrations at pH 6 in 20 mM histidine buffer. IgG concentrations in (B) and (C) range from 74.2 to 82.2 mg/mL. Different buffer systems were used in (B) and (C) to align with the experimental design; buffers in (B) were selected to match the target pH.

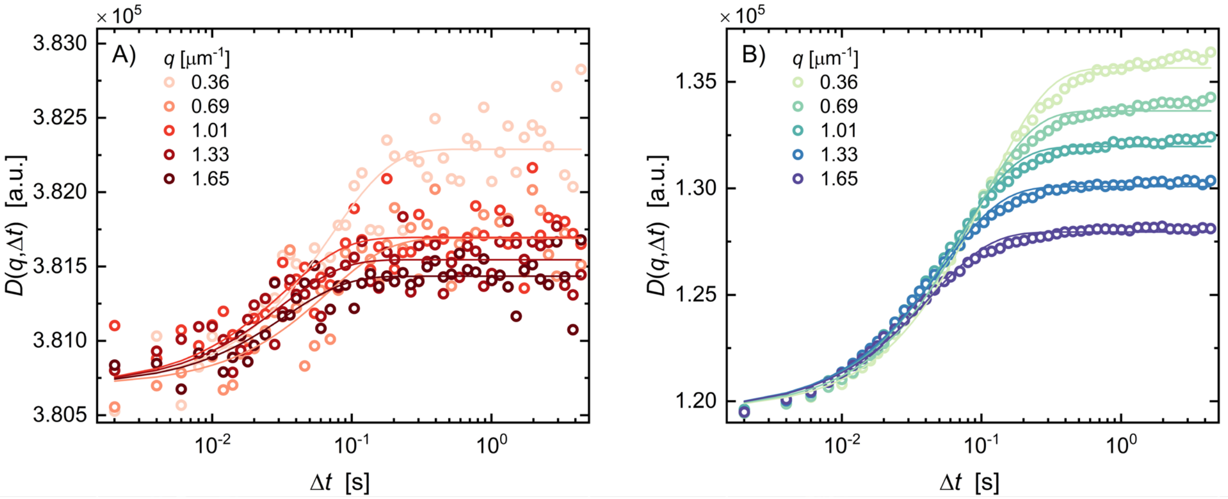


**Figure S2.** Representative image structure functions $D(q, \Delta t)$ obtained from (A) bright-field DDM and (B) phase-contrast DDM for 37.1 mg/mL IgG in MES buffer at pH 6, shown for five different scattering wavevectors $q$. Experimental data (open circles) are fitted to a single-exponential decay based on Eq. 2. (solid lines).

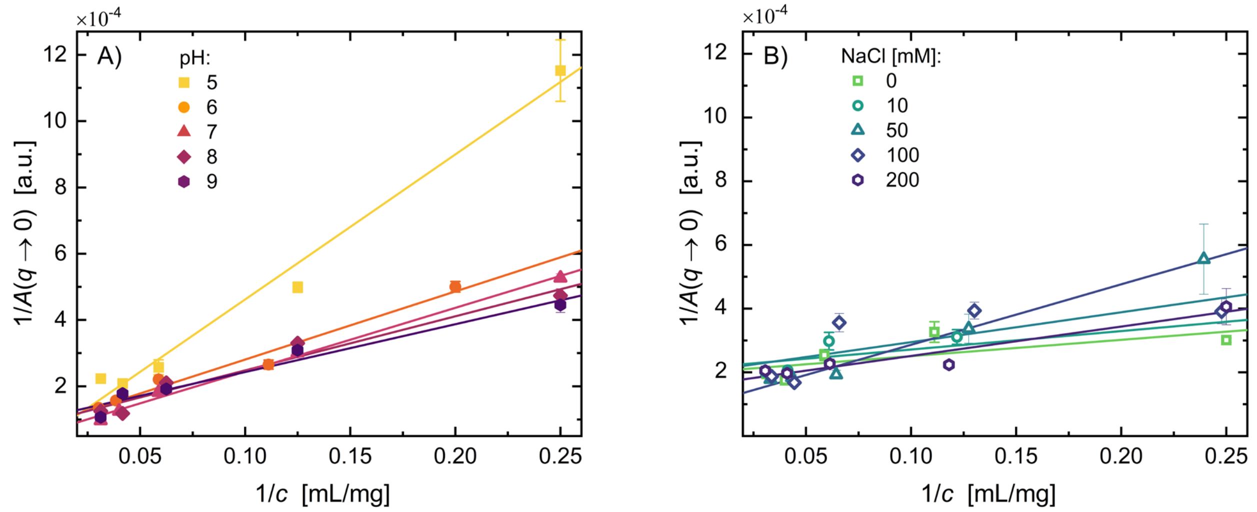


**Figure S3.** Linearized plot of DDM amplitude $A(q \to 0)$ versus protein concentration $c$ for IgG under (A) varying pH conditions with 100 mM added NaCl (solid symbols) and (B) varying NaCl concentrations at pH 6 in 20 mM histidine buffer (open symbols). Different buffer systems were used in (A) and (B) to align with the experimental design; buffers in (A) were selected to match the target pH. Error bars reflect the standard errors from the linear regression used to extrapolate $A(q)$ to $q \to 0$. The solid lines represent weighted fits based on Eq. 5.

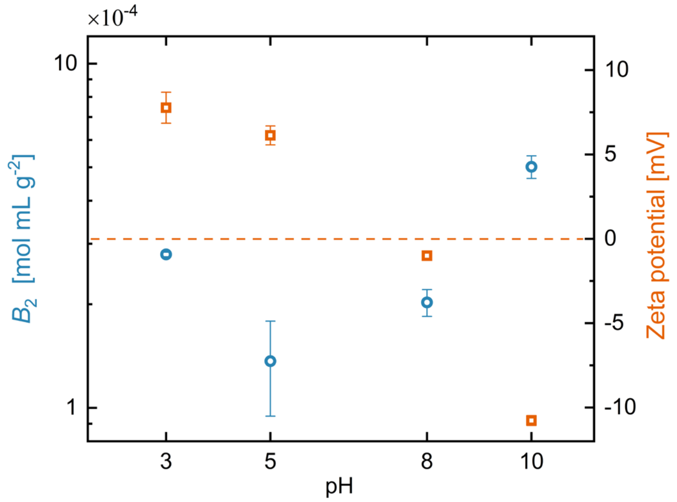


**Figure S4.** Second virial coefficient $B_2$ (blue circles) and zeta potential (orange squares) as a function of sample pH without added salt. The orange dashed line indicates the point of zero charge (PZC), where the zeta potential is zero and the net surface charge on IgG is neutral. Error bars on $B_2$ reflect standard errors from the weighted fits to Eq. 5, while those on the zeta potential indicate the standard error of the mean from three replicate measurements.

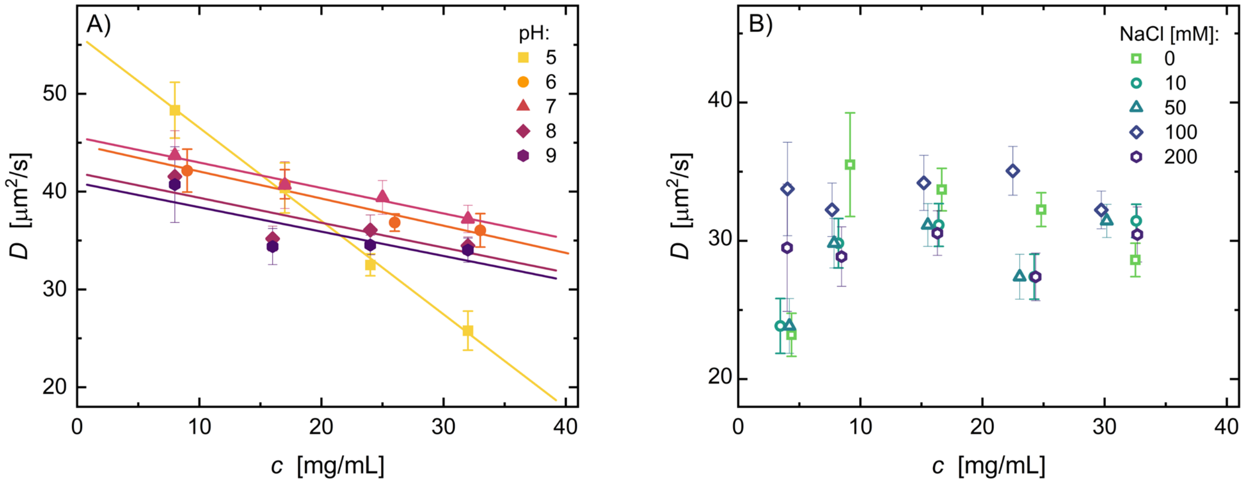


**Figure S5**. Diffusion coefficient $D$ (solid symbols) as a function of IgG concentration $c$ for (A) varying pH conditions with 100 mM added NaCl (solid symbols) and (B) varying NaCl concentrations at pH 6 in 20 mM histidine buffer (open symbols). Different buffer systems were used in (A) and (B) to align with the experimental design; buffers in (A) were selected to match the target pH. Error bars on $D$ correspond to standard errors determined from the weighted fits to Eq. 3. The solid lines in (A) show weighted fits based on Eq. 6, from which the diffusion interaction parameter $k_\mathrm{d}$ was obtained.

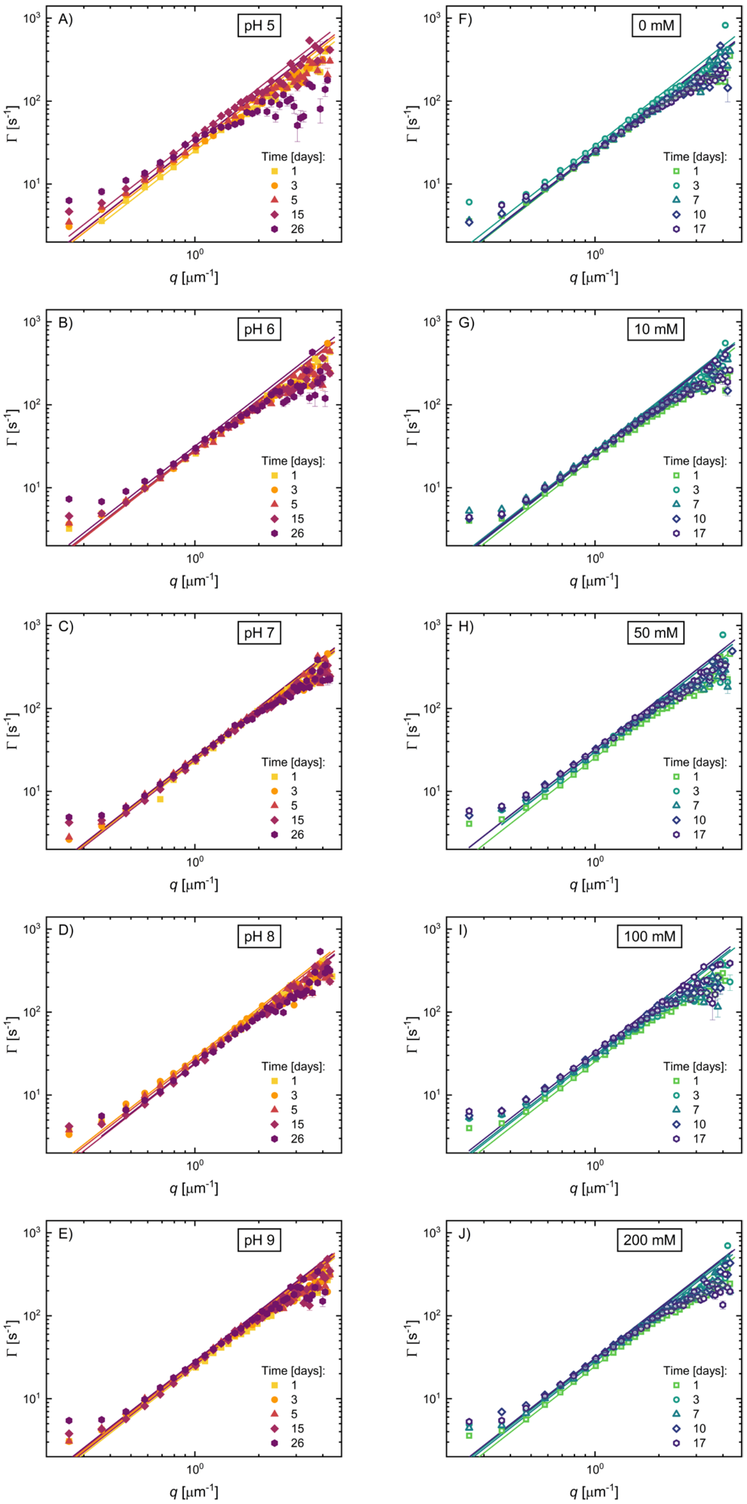


**Figure S6**. Relaxation rate $\Gamma(q)$ as a function of scattering wavevector $q$, with weighted fits to Eq. 3 (solid lines). Data are shown for five time points corresponding to those used to determine the diffusion coefficient $D$, and subsequently the hydrodynamic radius

$R_h$ in Figure 8. Panels (A) to (E) correspond to varying pH conditions with 100 mM NaCl (solid symbols), while panels (F) to (J) correspond to varying NaCl concentrations at pH 6 in 20 mM histidine buffer (open symbols). Error bars on $\Gamma(q)$ represent standard errors from the fits to Eq. 2 (some are smaller than the data markers).

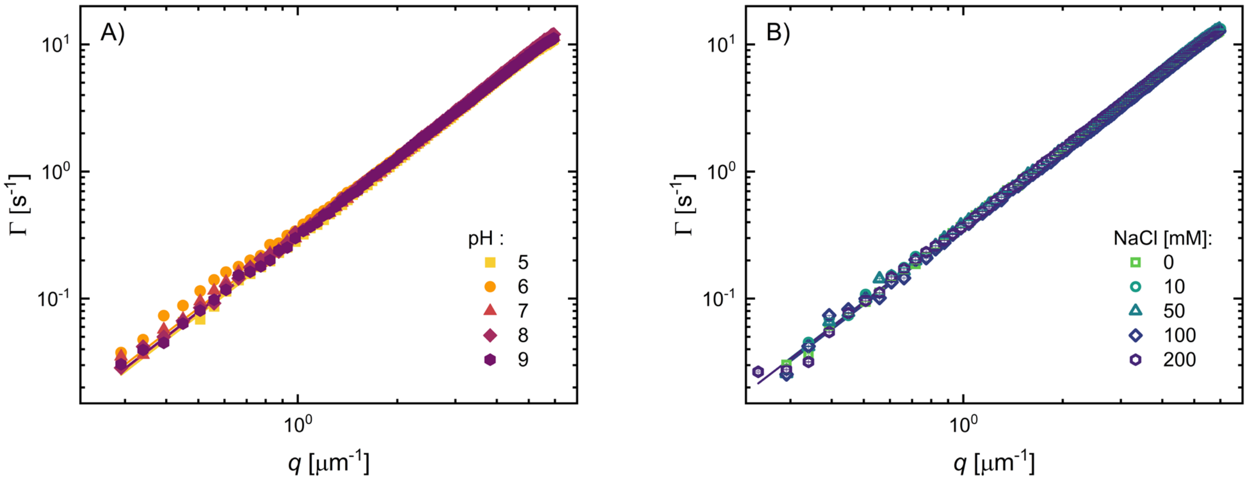


**Figure S7.** Relaxation rate $\Gamma(q)$ as a function of scattering wavevector $q$ under (A) varying pH conditions with 100 mM added NaCl (solid symbols) and (B) varying NaCl concentrations at pH 6 in 20 mM histidine buffer (open symbols). The solid lines correspond to weighted fits to Eq. 3 that were used to obtain the diffusion coefficient $D$, and subsequently the sample viscosity $\eta$ in Figure 9. Error bars on $\Gamma(q)$ represent standard errors from the fits to Eq. 2 (some are smaller than the data markers).
